# Dose-sparing self-amplifying RNA vaccine induces high functional antibodies to blood-stage *Plasmodium falciparum* malaria

**DOI:** 10.64898/2026.04.29.721807

**Authors:** Adam Thomas, Timothy Ho, Sandra Chishimba, Liriye Kurtovic, Claretta S. D’Souza, James Beeson

**Author notes:** <u>Corresponding authors:</u> James Beeson/Adam Thomas, Burnet Institute, 85 Commercial Road, Melbourne, VIC, 3004, Australia.

## Abstract

**Introduction:** Next-generation malaria vaccines are urgently needed to provide greater efficacy and longevity. Antibodies targeting blood-stage merozoites can confer protection against clinical malaria through multiple Fc-mediated functions. In particular, merozoite surface protein 2 (*Pf*MSP2), is a known target of protective antibodies that can clear merozoites via multiple antibody Fc-mediated functions, making is a highly promising vaccine candidate.

**Methods:** We developed *Pf*MSP2 as a self-amplifying RNA (saRNA) vaccine, which was successfully validated for *in vitro* expression in mammalian cells. Subsequently, the *Pf*MSP2-saRNA was formulated as lipid nanoparticles (LNP) and evaluated for immunogenicity in mice in a 3-dose regimen comparing 1 μg and 10 μg doses. We evaluated the induction of antibodies with functional activities relevant to protective immunity.

**Results:** Our *Pf*MSP2-saRNA vaccine induced antigen-specific IgG responses that recognised the surface of whole merozoites. Both 1 μg and 10 μg dosing induced comparable antibodies to *Pf*MSP2, and responses were predominantly murine cytophilic IgG subclasses. These vaccine-induced antibodies demonstrated potent Fc-mediated functions, including complement fixation and binding of human Fcγ-receptor I (FcγRI), after only two doses, which remained consistent after the third dose.

**Conclusions:** *Pf*MSP2 is highly immunogenic using the saRNA vaccine platform in a dose-sparing regimen, and induces antibodies with multiple Fc-mediated functions associated with protective immunity in humans. This saRNA platform is a promising strategy to develop highly efficacious vaccines requiring lower and fewer doses.

## Introduction

Malaria has a high global health burden, and the World Health Organisation (WHO) estimated more than 0.25 billion recorded cases and 0.6 million deaths in 2024 [1]. The highest disease morbidity is in Africa, with the majority of mortality being in children under 5 years. *Plasmodium falciparum* is the parasite species with high mortality globally, and there are risks of high resurgence in coming years. Therefore, effective vaccines are needed for malaria control and elimination [1].

*P. falciparum* has a complex lifecycle, whereby *Anopheles* mosquitoes transmit the sporozoite parasite through skin bites. The sporozoites infect hepatocytes in the liver and develop into merozoites, which are then released into the blood and undergo exponential asexual replication within red blood cells. This results in clinical malaria symptoms and can lead to severe illness and death [2, 3]. There are only two approved vaccines for malaria, RTS.S/AS01 and R21/MatrixM, that target the sporozoite to prevent initial hepatic infection [4, 5]. However, these vaccines provide only modest efficacy with breakthrough blood-stage parasitemia and clinical malaria still occurring, requiring booster doses annually [4]. Therefore, next-generation vaccines are needed to reduce malaria morbidity and mortality rates, and achieve greater efficacy and longevity [6]. Blood-stage merozoite antigens are attractive vaccine targets as they have the potential to reduce blood-stage replication and associated malaria symptoms. Antibodies targeting merozoites can inhibit blood-stage replication via Fc-mediated antibody functions including complement fixation and interaction with Fc receptors to promote opsonic phagocytosis and cellular cytotoxicity [7-10].

The *P. falciparum* merozoite surface protein 2 (*Pf*MSP2) is an abundant surface protein and potential vaccine candidate for blood-stage malaria [11-13]. *Pf*MSP2 has a variable region and dimorphic region comprising two allelic families (3D7 and FC27), and conserved N- and C-terminal domains [14]. Naturally acquired antibodies to *Pf*MSP2 have been associated with protection against malaria across different populations [12, 15]. These antibodies function via the antibody Fc region to mediate complement fixation and opsonic phagocytosis, leading to the inhibition of merozoites replication and clearance [7, 12, 16]. Therefore, inducing these protective antibody functions is important for future blood-stage vaccines.

Advances in vaccine technologies and the SARS -CoV2 mRNA vaccines’ success have introduced a new era of rapid vaccine development for infectious diseases [17]. The mRNA technology offers efficient single- and multi-antigen formulations, and rapid adaptability to emerging pathogens and strains [18]. Additionally, mRNA-lipid nanoparticle (LNP) vaccines induce robust antibody responses but there are limitations with standard non-replicating linear mRNA [19, 20]. Challenges include the requirement for high dosage, multi-dose regimens, and inefficient endosomal escape of mRNA [19, 21]. To overcome these limitations, a next-generation self-amplifying RNA (saRNA) platform can be utilised [22-24]. saRNA vectors include a viral replicase derived from an alphavirus that enables self-limiting replication of RNA transcripts *in situ* [25]. This feature of saRNA vaccines offers key advantages such as lower (and fewer) doses to elicit robust immune responses and antibody durability [24, 26]. A recent saRNA vaccine for SARS -CoV2 was approved in Japan and demonstrated a durable immune response compared to conventional mRNA vaccines, despite using a 6-20 fold lower dose [26, 27].

Here we designed and evaluated a *Pf*MSP2 vaccine using two dosing regimens of the saRNA-LNP platform with the aim of generating functional antibodies against merozoites. This work provides important proof-of-concept for the potential dose-sparing and shorter-regimen next-generation saRNA vaccines for malaria.

## Results

### PfMSP2 saRNA vaccine shows high immunogenicity in mice

We designed a near full-length *Pf*MSP2 (3D7 allele) saRNA vaccine based on a previously described *Pf*MSP2 vaccine construct with the addition of a C-terminal Flag-tag [11, 16]. The saRNA was codon-optimised and synthesised using an alphavirus vector. We first transfected Expi293F (HEK293) suspension cells with naked saRNA to confirm antigen expression. *Pf*MSP2 3D7 saRNA showed protein expression over time by detection of the C-terminal Flag-tag (Figure 1A). We then formulated the saRNA-LNP for *in vivo* testing in female C57BL/6 mice (n=5 per group) which received a fixed low-dose (3 × 1 μg) or high-dose (3 × 10 μg) on days 0, 28 and 56 (Figure 1B). Mouse sera were collected at baseline (pre-bleed) and after each vaccine dose, with the peak response time-point collected on day 70 (3^rd^ bleed). Vaccine-induced IgG to *Pf*MSP2 increased after the 2^nd^ dose in both groups, but interestingly, there was no significant increase after the 3^rd^ dose (Figure 1C). This indicates a dose-sparing advantage of the saRNA-LNP vaccine with fewer doses to achieve maximal IgG induction. We found that vaccine-induced antibodies can recognise native *Pf*MSP2 on the merozoite surface. ELISA was performed using whole merozoites that were isolated from *P. falciparum* blood-stage culture. Vaccine-induced IgG from the high-dose regimen at peak (day 70) was observed to bind native *Pf*MSP2 on the merozoite surface (Figure 1D). There was no significant difference between the saRNA and the recombinant *Pf*MSP2 vaccine control; however, the saRNA showed reduced variation between mice [16, 28].

**Figure 1.**
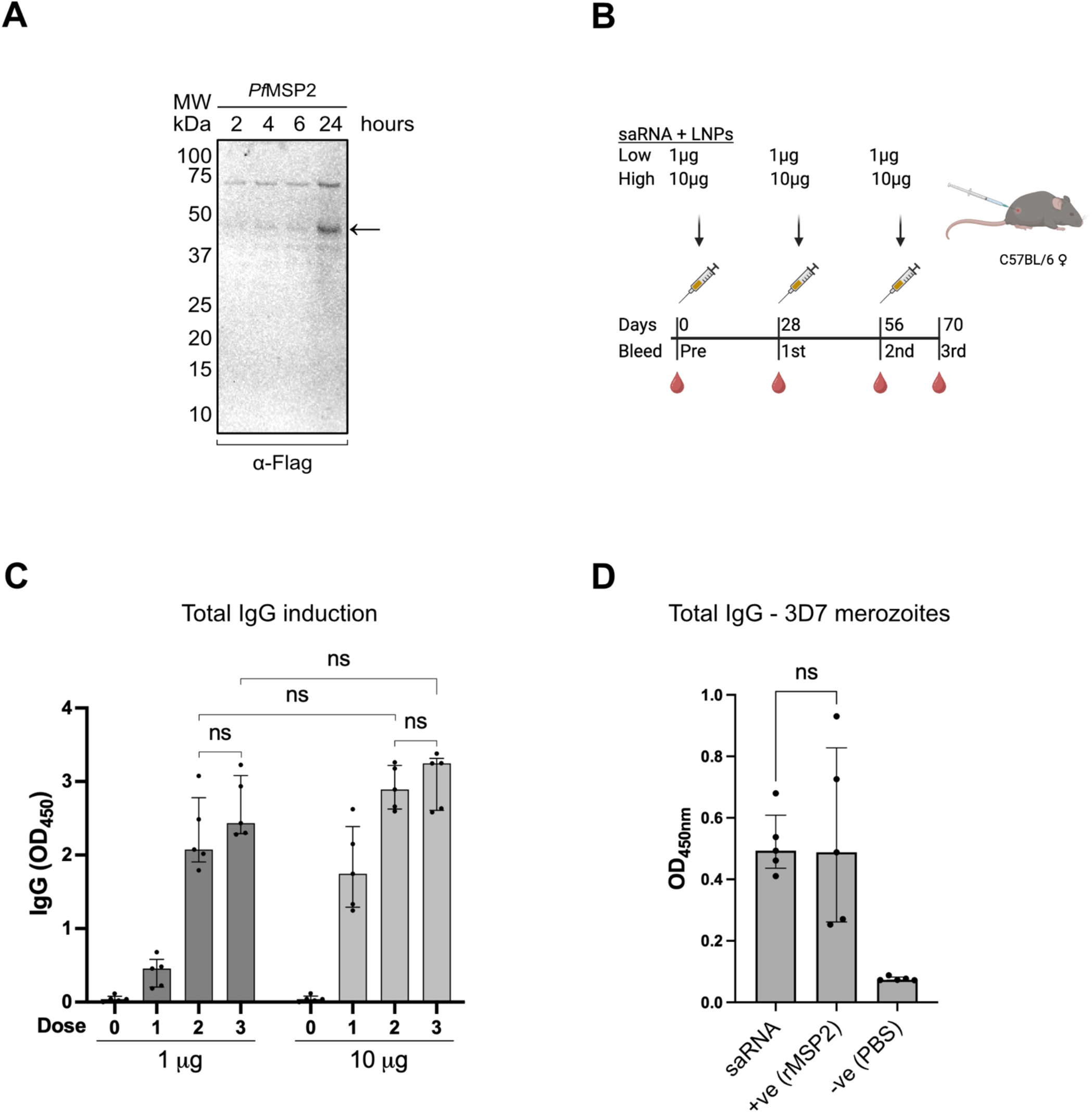
*Pf*MSP2 saRNA vaccine evaluation. (A) Expi293F cells were transfected with naked saRNA encoding *Pf*MSP2 (3D7 allele) and whole cells were collected. The samples were reduced for western blot, and protein expression was detected using α-Flag DyLight 800 monoclonal IgG (1/1000 dilution). Arrow indicates *Pf*MSP2 band. (B) Schematic of mouse immunisation regimens and timeline for blood collection (low dose: 3 × 1 μg; high dose: 3 × 10 μg saRNA *Pf*MSP2). Mouse bleeds were also collected prior to immunisation. (C) IgG reactivity in mice after immunisation with saRNA-LNP *Pf*MSP2. Mouse sera measured at 1/3200 dilution by ELISA (n=5 per group). Assays were performed in duplicate and error bars represent median ± IQR. Statistical analysis was performed using an unpaired nonparametric Mann-Whitney test; ns indicates non-significance (p>0.05). (D) ELISA analysis of IgG antibody reactivity to native *Pf*MSP2 on whole merozoites using mouse sera from saRNA 10 μg (high-dose) vaccination on day 70 at 1/100 dilution (n=5 per group), and control recombinant *Pf*MSP2 vaccine-induced antibodies on day 70 at 1/100 dilution (n=5 per group) [16]. Error bars represent median ± 95% CI and assays performed in duplicates. Statistical analysis was performed using an unpaired nonparametric Mann-Whitney test; ns indicates non-significance (p>0.05).

### PfMSP2 saRNA vaccines induce high functional lgG with Fc-mediated activity

To understand the protective potential of immune responses, the profiles of mouse IgG subclasses induced by saRNA *Pf*MSP2 vaccines were examined [29]. Remarkably, the murine IgG2b and IgG2c that confer Fc-mediated protective functions were strongly induced, with low induction of IgG1 (which has low Fc-mediated functional activity) in serum following vaccination [14, 16, 30, 31]. Testing IgG after each dose, the two dosing regimens showed comparable subclass profiles with IgG2c and IgG2b at higher magnitude, but low IgG1 and IgG3 levels (Figure 2A-B). The subclass levels only moderately increased after the 3^rd^ dose in both regimens, in line with the total IgG results. Previous studies have shown that antibodies to *Pf*MSP2 have Fc-mediated functions that include complement fixation and activation, as well as Antibody-Dependant Cellular Inhibition (ADCI) and opsonic phagocytosis of merozoites that are mediated by IgG interactions with Fcγ receptors [7, 11, 15, 29, 32-34]. Therefore, we examined Fc-mediated functions of saRNA vaccine-induced mouse antibodies. Interestingly, the murine IgG demonstrated strong C1q fixation, which is the first step of classical complement activation (Figure 2C). Vaccine-induced antibodies also showed strong binding to Fcγ Receptor I, which can promote phagocytosis and other functions (Figure 2D). All functional activities were broadly comparable between groups, with the 10 μg saRNA regimen showing significantly higher activity after the 2^nd^ dose, albeit not 10-fold higher than the 1 μg saRNA regimen. No significant difference was observed after the 3^rd^ dose between the two regimens, indicating a dose-sparing potential and fewer doses required of the saRNA vaccine.

**Figure 2.**
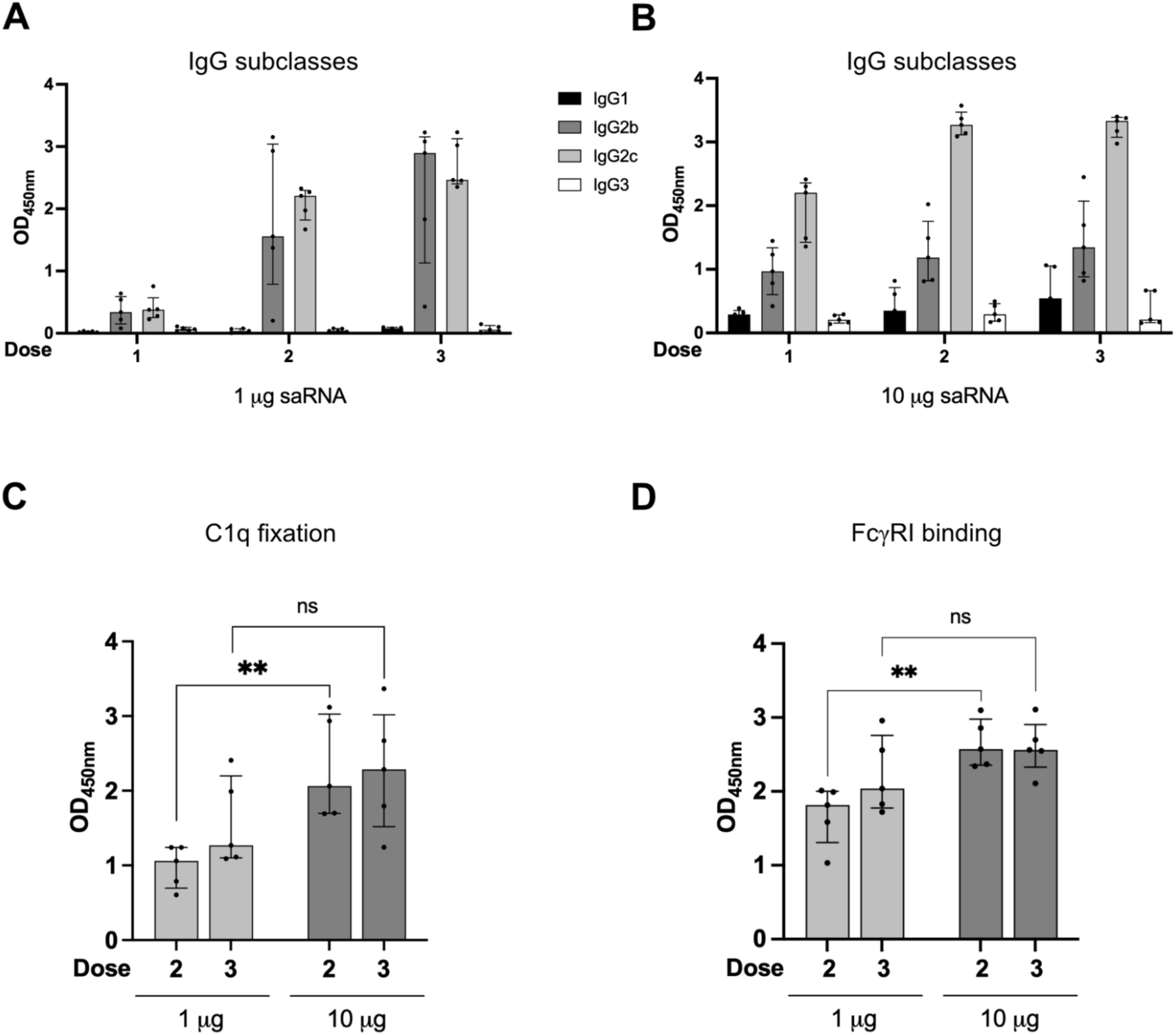
Functional profile of *Pf*MSP2 saRNA vaccine-induced IgG. (A) ELISA analysis of mouse IgG subclasses in sera (n=5 per group) of 1 μg saRNA regimen. Assays were performed in duplicates (1/3200 dilution) and error bars represent median ± IQR. (B) Mouse IgG subclasses in sera (n=5 per group) of 10 μg saRNA vaccinated mice. Assays were performed in duplicates (1/3200 dilution) and error bars represent median ± IQR. (C) Human C1q fixation by murine IgG from saRNA vaccine regimens (1/400 dilution). Assays were performed in duplicates and error bars represent median ± IQR; Statistical analysis was performed using an unpaired nonparametric Mann-Whitney test; ** is significant (p=0.0079), ns indicates non-significance (p>0.05). (D) Human FcγRI binding (1/12800 dilution) by murine IgG from sera. Assays were performed in duplicates and error bars represent median ± IQR; Statistical analysis was performed using an unpaired nonparametric Mann-Whitney test; ** is significant (p=0.0079), ns indicates non-significance significant (p>0.05).

## Discussion

In this study we demonstrate a new strategy to target the merozoite surface using saRNA vaccines to generate antibodies with Fc-mediated functional activity. We designed the saRNA vaccine against *Pf*MSP2 because it is an abundant merozoite surface antigen and a known target of functional antibodies that promote complement activation to inhibit blood-stage replication and mediate opsonic phagocytosis of merozoites [7, 12, 32, 34]. Furthermore, antibodies to *Pf*MSP2 have been associated with protection in studies of naturally acquired and vaccine-induced immunity [12, 32]. We found the vaccine effectively generated antibodies with Fc-mediated functional activities using a dose-sparing 1 μg regimen with two doses.

We validated the expression of *Pf*MSP2 in mammalian cells before progressing the saRNA vaccine into preclinical animal studies. The saRNA-LNP formulation induced optimal levels of cytophilic IgG subclasses in mice following two doses for either 1 μg or 10 μg regimens, unlike conventional mRNA vaccines that require three doses [16]. The efficiency of conventional mRNA vaccine translation *in vivo* has been reported to be 1-2% due to endosomal escape limitations and requires higher and repeated mRNA doses [20, 21, 35]. In contrast, saRNA offers self-replication *in situ* allowing for prolonged and increased antigen presentation with a lower dose, thus decreasing the impact of low endosomal escape [26, 27, 36, 37]. Remarkably, the immunogenicity of the saRNA vaccine was comparable between both dosing regimens (1 μg and 10 μg), with reduced biological variation between mice compared to a recombinant protein vaccine [16]. There was no clear improvement in immunogenicity using higher or more frequent doses of saRNA demonstrating dose-sparing advantages. However, further dose optimisation studies and larger sample sizes would be needed to more formally evaluate these results.

saRNA vaccine-induced antibodies showed strong functional activity, including complement fixation and interaction with FcγRI [38, 39]. This functional antibody profile is similar to what has been reported for antibodies in naturally acquired immunity, clinical trials of a recombinant *Pf*MSP2 protein vaccine, and *Pf*MSP2 conventional mRNA vaccine preclinical studies [7, 12, 16]. Fcγ receptor engagement and complement activation functions have been shown to be key to parasite inhibition and clearance *in vitro* and *in vivo* [7, 12, 34, 40]. The IgG2c and IgG2b subclasses of C57BL/6 mice, which can mediate antibody functions effectively, were nearly exclusively induced using the saRNA platform [42]. Thus, it was promising that the 1 μg dose-sparing saRNA-LNP regimen showed similar induction of antibodies to the 10 μg regimen. This is highlighted by the 10-fold lower dose that did not reflect a 10-fold decrease in antibody magnitude or function. Naturally acquired antibodies reportedly target the variable allele-specific regions of *Pf*MSP2 [12, 15]. Therefore, further construct optimisation and inclusion of the FC27 allele are needed in future studies to broaden protection [14, 28, 41]. Additionally, understanding cellular immune responses may provide insights into vaccine longevity and potential protection. Due to the absence of *Pf*MSP2 or an orthologue in rodent malaria species, and differences in C1q and Fc-mediated functions between humans and mice, laboratory mouse models pose limitations for assessing the efficacy of *Pf*MSP2-based vaccines [42]. Therefore, this saRNA vaccine could be further evaluated in non-human primate studies or controlled human malaria infection (CHMI) vaccine trials.

## Conclusions

Our study provides a proof-of-concept for using saRNA as a platform for malaria vaccines, demonstrating strong immunogenicity and antibody functional activities. We showed that the saRNA platform can be used in dose-sparing and fewer dose regimens that induce functional IgG against *Pf*MSP2. A low-dose and 2-dose regimen induced strong immunity comparable to the high-dose regimen. This platform can be extended to include other malaria vaccine antigens in development, such as *Pf*Rh5, to induce antibodies with both growth-inhibitory and Fc-mediated functional activities. This platform could offer greater protective vaccine efficacy with short and low dose vaccine regimens to achieve high-level protection against malaria.

## Materials and Methods

### Production of naked saRNA and saRNA-LNP vaccines

The codon-optimised *P. falciparum* MSP2 3D7 sequence (GenBank accession number JN248383 with the substitution of the C-terminal poly-Histidine tag with a Flag-tag) was synthesised as described previously by GenScript Biotech [25] using saRNA-2 vector with 5’ Cap1, 3’ Poly(A) sequence, and 5-methylcytidine substitution. The saRNA vaccine in this study used Moderna’s LNP formulation (SM102) in 1 mM Sodium citrate buffer (pH 6.5). All saRNA synthesis, LNP formulation, and characterisation were performed by GenScript Biotech.

### Expression and validation of naked mRNA

The expression of naked saRNA was performed in Expi293F suspension cells (Gibco) using TransIT-mRNA (Mirus) transfection kit and analysed via NuPAGE and western blot as described previously [16].

### PfMSP2 production

Recombinant *Pf*MSP2 3D7 was manufactured at GroPep Pty Ltd (Adelaide, Australia) as previously described [11, 16].

### Study design

he evaluation of the *Pf*MSP2 saRNA-LNP vaccine was conducted using female C57BL/6 mice aged 8-12 weeks (n=5 per group). The mice were immunised with two regimens of saRNA-LNP: a fixed low-dose regimen (3 × 1 μg) and a fixed high-dose regimen (3 × 10 μg). The mice received 3 doses at 28 days intervals with prior blood collection preceding the injection. The immunisation and blood collection schedules are shown in Figure 1B. Mice received all immunisations via intramuscular injections to the calf muscle. The first two collections are retro-orbital bleeds, the subsequent collections are mandible bleeds, and the terminal collection is a cardiac bleed (following inhaled anaesthetic euthanasia). All mice were housed, cared for, and handled (injection and bleeding) by the WEHI Antibody Facility in Bundoora (Melbourne, Australia).

### Culture and isolation of P. falciparum merozoites

Parasite culture (*Plasmodium falciparum* 3D7) and merozoite isolation were performed as previously described [34].

### Expression analysis

To assess the expression of *Pf*MSP2 saRNA, the transfection samples were run on a gel under reducing conditions as previously described [16]. Briefly, whole-cell suspensions were prepared with 100 mM dithiothreitol (Thermo Fisher Scientific), 1 × NuPAGE LDS Sample Buffer (Invitrogen), and boiled at 95°C for 5 minutes, then vortexed for 30 seconds. Samples were loaded onto a 4-12% Bis-Tris gel (Invitrogen) with Precision Plus Protein Dual Color Standards (Bio-Rad) and run at 135 V and 200 mA for 1.5 hours in 1 × MES running buffer (Invitrogen).

Western blot analysis was performed as described previously [16]. The gel was transferred onto a nitrocellulose (NC) membrane using iBlot 2 Transfer Stacks (Thermo Fisher Scientific), then blocked with 1% (w/v) Bovine Serum Albumin (BSA; Bovogen) in 0.05% (v/v) Tween20 (Sigma-Aldrich) in 1 × PBS (PBS-T) overnight at 4 °C. The membrane was probed with mouse anti-Flag DyLight 800 (1/1000; Invitrogen) in 1-2% (w/v) BSA at RT with extensive washing with PBS-T between steps. The membrane was then visualised using the ChemiDoc MP (Bio-Rad).

### ELISA and plate-based assays

ELISA protocols were performed as described previously for IgG, C1q fixation, and human Fc receptor binding [12, 16, 29, 32, 40, 43, 44]. Assays were normalised across plates for individual experiments.

### Statistical analysis

All statistical analyses and graph generation for this study were conducted using GraphPad Prism 11 (MacOS) software. Unpaired nonparametric Mann-Whitney test was used to compare two groups.

## Acknowledgements

We thank Paul Masendycz, Ridouan Bouhbouh, and Myha Huynh at WEHI Antibody Facility for their support with our animal studies. Schematics were created using BioRender. Burnet Institute is located on the traditional lands of the Boonwurrung people of the Kulin nation.

## Ethics

Burnet Institute malaria antigens saRNA handling approval (IBC 48659). WEHI Antibody Facility animal in Bundoora Scientific Procedure Premises Licence (SPPL201904), and ethics approval (2020.019 and 2023.012).

## Author contribution

AT and JB designed the study. AT, TH, and SC conducted the experiments. AT and TH were involved in data analysis with guidance from LK, SC, CD, and JB. AT led the writing of the manuscript with guidance from all authors. All authors reviewed and approved the final version.

